# NusG links transcription and translation in *Escherichia coli* extracts

**DOI:** 10.1101/2021.07.31.454578

**Authors:** Elizabeth J. Bailey, Max E. Gottesman, Ruben L. Gonzalez

## Abstract

In bacteria, transcription is coupled to, and can be regulated by, translation. Although recent structural studies suggest that the N-utilization substance G (NusG) transcription factor can serve as a direct, physical link between the transcribing RNA polymerase (RNAP) and the lead ribosome, mechanistic studies investigating the potential role of NusG in mediating transcription-translation coupling are lacking. Here, we report development of a cellular extract- and reporter gene-based, *in vitro* biochemical system that supports transcription-translation coupling as well as the use of this system to study the role of NusG in coupling. Our findings show that NusG is required for coupling and that the enhanced gene expression that results from coupling is dependent on the ability of NusG to directly interact with the lead ribosome. Moreover, we provide strong evidence that NusG-dependent coupling enhances gene expression through a mechanism in which the lead ribosome that is tethered to the RNAP by NusG suppresses spontaneous backtracking of the RNAP on its DNA template that would otherwise inhibit transcription.

## Introduction

In 2010 Proshkin *et al.* showed not only that the rates of transcription and translation match in bacteria, as was widely accepted, but that the rate of translation influenced the rate of transcription [1]. Conceptually, this is fascinating, as it suggests that transcription and translation do not proceed in a purely sequential manner, but rather that there is regulatory communication, of some sort, between the two processes such that the second process, translation, has control over the first process, transcription.

Contemporaneously with Proshkin *et al.*’s report, Burmann *et al.* proposed a structure-based, molecular mechanism for the possible communication between transcription and translation (*i.e.*, transcription-translation coupling) [2]. Specifically, they presented structural evidence that the carboxy (C)-terminal domain (CTD) of the 21 kDa N-utilization substance (Nus) G transcription factor can physically interact with ribosomal protein uS10. Moreover, the surface of uS10 with which the NusG CTD was shown to interact is solvent exposed and available within the context of the ribosomal small, or 30S, subunit, suggesting that the NusG CTD can directly interact with the 30S subunit and an intact, translating 70S ribosome. Notably, a flexible, 15-amino acid linker connects the NusG CTD to the amino (N)-terminal domain (NTD) of NusG, a domain that was already known to directly interact with RNA polymerase (RNAP) [3]. Taken together, these data suggested a model in which NusG could simultaneously bind the transcribing RNAP and the translating lead ribosome, physically linking the two processes.

Physical tethering of RNAP to the ribosome by NusG is further supported by biochemical and structural studies of NusG binding to the 70S ribosome [4, 5], as well as to both RNAP and the 70S ribosome within the context of transcription-translation complexes [4, 6–8]. Structural analysis of transcription-translation complexes assembled from purified components on relatively long mRNAs in which NusG is observed to tether RNAP to the 70S ribosome show that the NusG NTD contacts the β’ and β subunits of RNAP, while the NusG CTD contacts the solvent-accessible surface of uS10 within the 30S subunit of the 70S ribosome [6, 7]. One of the structures further suggests that the NusA transcription elongation factor participates in the complex [7], a finding that is supported by a recent, in-cell, cryogenic electron tomography study [8]. In structures of transcription-translation complexes analogously assembled on relatively shorter mRNAs, NusG is excluded from the complex and a direct link between RNAP and the 70S ribosome is observed [6, 7]. Notably, the relative orientation of RNAP and the 70S ribosome observed in the structure of an RNAP-70S ribosome complex formed by colliding a translating 70S ribosome into a stalled RNAP in the absence of NusG [9] is consistent with that observed in the structures of transcription-translation complexes assembled on relatively short mRNAs [6, 7]. In contrast, the relative orientation of RNAP and the 70S ribosome inferred from biochemical experiments [10] and observed in a structure [11] of RNAP-70S ribosome complexes assembled in the absence of either mRNA and NusG is inconsistent with that observed for any of the transcription-translation complexes [4, 6–8].

As the previous paragraph demonstrates, the recent, spectacular progress in our understanding of the structural basis of transcription-translation coupling has generated a number of compelling, structure-based mechanistic hypotheses regarding the mechanism of transcription-translation coupling and its role in regulating gene expression. Unfortunately, however, a paucity in the availability of *in vitro* experimental systems allowing full biochemical control over the factors that mediate transcription-translation coupling has thus far limited comprehensive testing of these hypotheses. To address these technological and knowledge gaps, here we report the development of such an *in vitro* biochemical system and the use of this system to study the role of NusG in transcription-translation coupling. Specifically, we have used *Escherichia coli* S30 cellular extracts and a luciferase reporter gene construct to develop an *in vitro* biochemical system that preserves the coupling between transcription and translation. Addition of a DNA template encoding luciferase to the S30 extracts enables us to conduct transcription-translation reactions, whereas addition of a separately and independently *in vitro* transcribed mRNA encoding luciferase permits us to decouple translation from transcription and perform translation-only reactions. Importantly, this system allows us to control the presence, identities, and concentrations of factors mediating transcription-translation coupling, thereby allowing us to test structure-based hypotheses, investigate the mechanism of transcription-translation coupling, and elucidate the molecular consequences of uncoupling transcription from translation. Using this system in combination with wildtype and mutant variants of NusG and RNAP, we have investigated the role that NusG-mediated tethering of RNAP to the lead ribosome plays in transcription-translation coupling and the mechanism through which such tethering allows the rate of translation to influence the rate of transcription. The results we present here provide strong evidence supporting a mechanistic model in which tethering of RNAP to the lead ribosome by NusG increases the efficiency of gene expression through a mechanism in which the tethered lead ribosome suppresses backtracking of RNAP that would otherwise impair transcription.

We began our work by attempting to generate an *E. coli* S30 cellular extract that would completely lack endogenous NusG such that it could serve as a standard extract to which we could add exogenously overexpressed and purified NusG proteins and perform transcription-translation and translation-only reactions. We were motivated to generate such an S30 extract based on previously published *in vivo* cell biology studies showing that *E. coli* strains in which the gene encoding NusG, *nusG*, had been deleted are viable, albeit extremely slow growing [12, 13]. Following up on these previous studies, we performed *in vivo* cell biology experiments using a *nusG* deletion strain prepared in an *E. coli* MDS42 background (Supplementary Materials and Methods). Providing a rationale for the previously observed extremely slow growth phenotype [12, 13], this strain expressed 10-fold less β-galactosidase than a wildtype MDS42 strain, a defect that could be fully complemented by expression of a plasmid-borne copy of wildtype *nusG* (Supplementary Figure S1). Based on the ability of plasmid-based expression of NusG to complement the lack of endogenously expressed NusG, we generated an *E. coli* MG1655-based strain in which *nusG* had been deleted (MG1655 *ΔintR-kilR::Cam*^*R*^ *nusG::Kan*^*R*^), hereafter referred to as the *nusG* knock-out (KO) strain (Materials and Methods). Western blot analyses against the β’ subunit of RNAP and ribosomal protein S3 established that the RNAP and ribosome content of S30 extracts prepared from the *nusG* KO strain was similar to that of S30 extracts prepared from the wildtype MG1655 parent strain (MG1655 *ΔintR-kilR::Cam*^*R*^; Supplementary Materials and Methods and Supplementary Figure S2).

Using a circularized DNA plasmid encoding the firefly luciferase gene downstream from a *tac* promoter and a ribosome binding site (pBESTluc, Promega) as a template, we next measured the luminescence activity of the luciferase expressed in transcription-translation reactions performed in S30 extracts prepared from the *nusG* KO strain (Materials and Methods). Initial experiments resulted in luciferase activities that were 5-fold lower than analogous experiments performed in identically prepared S30 extracts from the wildtype MG1655 parent strain. Given our *in vivo* results (Supplementary Figure S1), we were surprised to observe that addition of purified wildtype NusG (NusG-WT) (a kind gift from Prof. Paul Röche, University of Bayreuth) to the reactions did not restore the luciferase activity (Supplementary Figure S3). Collectively, these results strongly suggest that *nusG* KO extract is deficient in luciferase expression and that this defect is irreversible in our *in vitro* transcription-translation system.

In an attempt to identify the molecular basis for the irreversible defect in luciferase expression that we observed in the *nusG* KO S30 extracts, we performed next-generation RNA sequencing (RNA-Seq) of the *nusG* KO strain and, as a reference, the wildtype MG1655 parent strain, in order to identify mRNAs whose cellular populations were up- or down-regulated upon deletion of *nusG* (Supplementary Materials and Methods). The results showed that the populations of a number of mRNAs encoding proteins with direct or indirect roles in translation were significantly deregulated in the *nusG* KO strain (Supplementary Tables S1 and S2). Thus, the *nusG* KO strain likely harbors pleiotropic defects in translation, explaining why simple addition of purified NusG-WT to the *nusG* KO extract could not rescue luciferase expression. Consistent with this interpretation, experiments in which an mRNA that had been *in vitro* transcribed from the pBESTluc plasmid was directly added to *in vitro* translation-only reactions performed in S30 extract prepared from the *nusG* KO strain showed that luciferase activity remained extremely low, confirming that S30 extracts prepared from the *nusG* KO strain were defective in translation (Supplementary Figure S4).

Given we could not generate a standard extract lacking NusG without introducing pleiotropic translation defects, we instead used the wildtype MG1655 parent strain to generate an S30 extract containing endogenous levels of NusG-WT, hereafter referred to simply as ‘standard extract’ (Materials and Methods). Addition of excess concentrations of purified NusG-WT or mutant NusG proteins to standard extract would then allow us to assess the effects of these proteins on the *in vitro* transcription-translation and translation-only activities of S30 extracts containing an endogenous level of NusG-WT (*i.e.*, how the added proteins modulate and/or compete with the endogenous level of NusG-WT).

To investigate the role of NusG in coupling translation to transcription, we first tested how addition of 1 μM of NusG-WT or each of two previously reported, purified mutant NusG proteins (a kind gift from Prof. Röche) to standard extract affected the luciferase activities of transcription-translation or translation-only reactions (Figure 1). The first mutant NusG protein is a truncation mutant in which the CTD has been deleted (NusG-NTD) such that the mutant protein is no longer capable of bridging RNAP and uS10 (5, 6). The second is a substitution mutant in which the phenylalanine at residue position 165 within the NusG CTD has been mutated to an alanine (NusG-F165A) (5, 6). Phenylalanine 165 is a NusG CTD residue that is highly conserved across bacteria [2, 5], forms part of its uS10-interacting surface (5, 6), and whose mutation to alanine we have previously shown disrupts the interaction of the NusG CTD with uS10 [2]. Notably, we have previously reported nuclear magnetic resonance (NMR) spectroscopy [2, 14] and *in vitro* transcription [5] studies demonstrating that these NusG-WT, NusG-NTD, and NusG-F165A proteins are properly folded and exhibit the expected biochemical activities. Consistent with this, we have also reported cell biology studies suggesting that NusG-NTD, and, by extension, NusG-F165A, compete with NusG-WT for binding to RNAP [3] (Materials and Methods).

**Figure 1.**
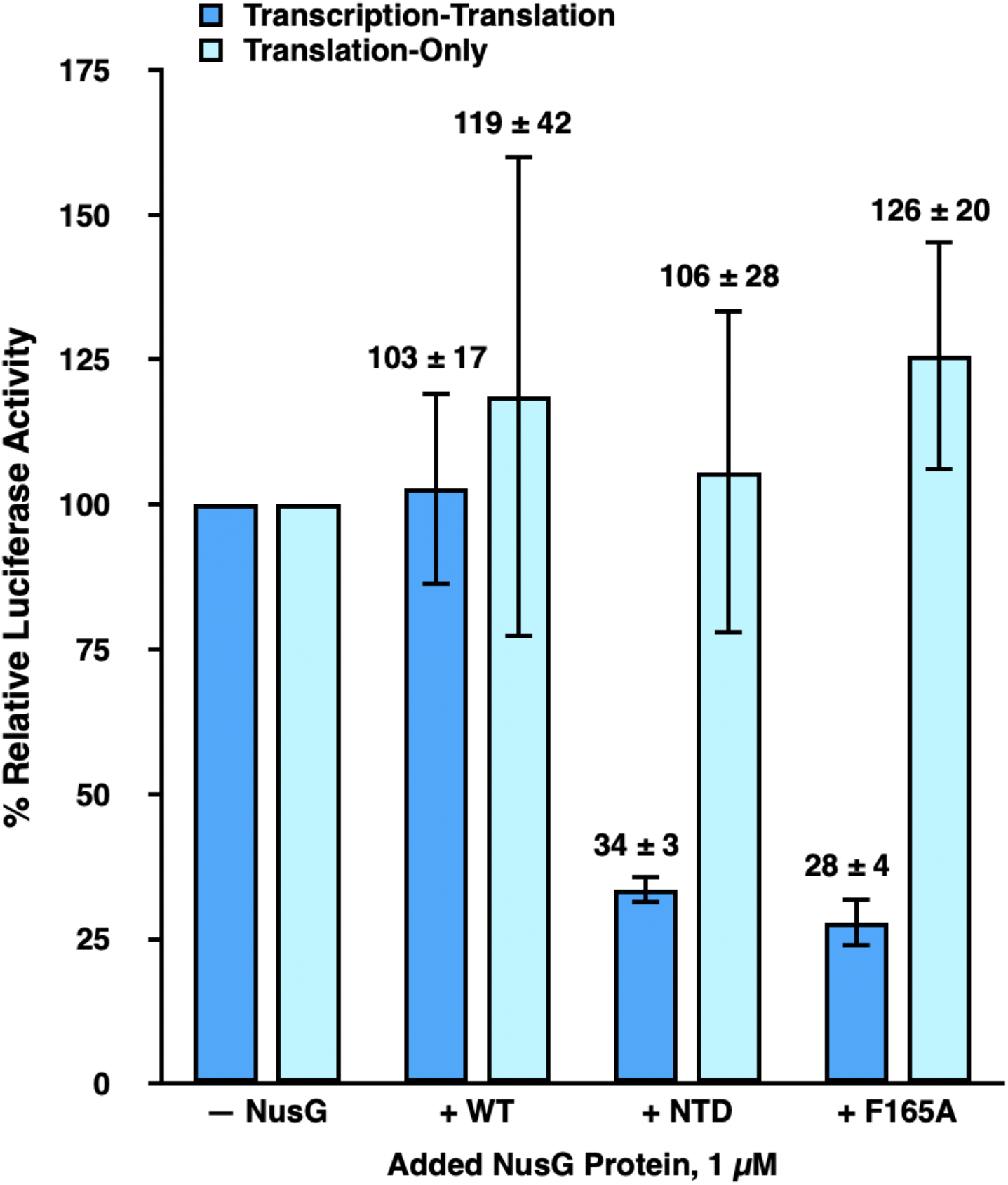
NusG-mediated tethering of RNAP to the lead ribosome enhances gene expression in transcription-translation reactions. Bar graph plotting the % relative luciferase activity of transcription-translation (dark blue bars) and translation-only (light blue bars) reactions performed using standard extract in the absence of any added NusG proteins (− NusG) or in the presence of 1 μM added NusG-WT (+ WT), NusG-NTD (+ NTD), and NusG-F165A (+ F165A). The luciferase activities of the transcription-translation and translation-only reactions in WT, NTD, and F165A are reported relative to those of the transcription-translation or translation-only reactions, respectively, in – NusG, which are each set to 100%. Error bars represent the standard deviations of each measurement.

To analyze the results of these experiments, we calculated the % relative luciferase activity of each transcription-translation or translation-only reaction, given as [(luciferase activity of the reaction performed in standard extract in the absence of added NusG proteins or in the presence of NusG-WT, NusG-NTD, or NusG-F165A proteins, as designated) / (luciferase activity of a corresponding ‘reference’ reaction analogously performed in standard extract in the absence of added NusG proteins) × 100] (Materials and Methods). Addition of NusG-WT to transcription-translation and translation-only reactions resulted in a % relative luciferase activity of 103 ± 17 % and 119 ± 42 %, respectively. These results suggest that addition of NusG-WT does not significantly alter the expression of luciferase in either transcription-translation or translation-only reactions performed in standard extract. In contrast, the % relative luciferase activities of added NusG-NTD or NusG-F165A in transcription-translation reactions were significantly decreased, to 34 ± 3 % and 28 ± 4 %, respectively, indicating that addition of NusG-NTD or NusG-F165A markedly reduces expression of luciferase in transcription-translation reactions performed in standard extract. Notably, the % relative luciferase activities of added NusG-NTD or NusG-F165A in translation-only reactions were 106 ± 28 % and 126 ± 20 %, respectively, suggesting that addition of NusG-NTD or NusG-F165A does not significantly alter translation of the luciferase-encoding mRNA.

The fact that added NusG-NTD and NusG-F165A decrease the expression of luciferase in transcription-translation reactions but have no effect on the expression of luciferase in translation-only reactions indicates that the interaction between the NusG CTD and uS10 within the ribosome, and, presumably, the attendant coupling of RNAP to the lead ribosome, is necessary for robust synthesis of the luciferase-encoding mRNA. Furthermore, assuming that NusG-F165A inhibits transcription through the same mechanism as NusG-NTD (*i.e.*, through abrogating the NusG CTD-uS10 interaction and RNAP-ribosome coupling), the fact that NusG-F165A exhibited as strong an inhibition as NusG-NTD confirms that the F165A substitution effectively disrupts the interaction of the NusG CTD with uS10 and therefore the coupling of RNAP to the lead ribosome in an *in vitro* cellular extract system. This is consistent with results from Saxena *et al.* demonstrating that the F165A substitution completely disrupted binding of NusG to ribosomes *in vitro* and significantly weakened the affinity of NusG for ribosomes *in vivo* [5]. The present results indicate that the F165A substitution mutation disrupts transcription as effectively as a full deletion of the NusG CTD.

There are two likely ways in which NusG-NTD and NusG-F165A could disrupt RNAP-ribosome coupling and inhibit transcription. The first is by allowing uncoupled RNAP to outpace the lead ribosome, generating naked mRNA that becomes available to the Rho transcription termination factor and permits premature, Rho-dependent transcription termination. The second is by allowing uncoupled RNAP to fall into long pauses and backtrack on the DNA template, effectively inhibiting transcription. To test for Rho-dependent termination, we asked if bicyclomycin (BCM), an antibiotic that selectively inhibits Rho [12, 15, 16], could rescue the inhibition of transcription induced by addition of 1 μM NusG-F165A to the standard extract used in transcription-translation reactions (Figure 2). The results of these experiments showed that titrating BCM over two orders of magnitude, from 0–700 μM, did not significantly restore luciferase activity in standard extract with 1 μM added NusG-F165A protein. Thus, premature transcription termination by Rho does not account for the failure to synthesize the luciferase mRNA in the absence of NusG-mediated RNAP-ribosome coupling.

**Figure 2.**
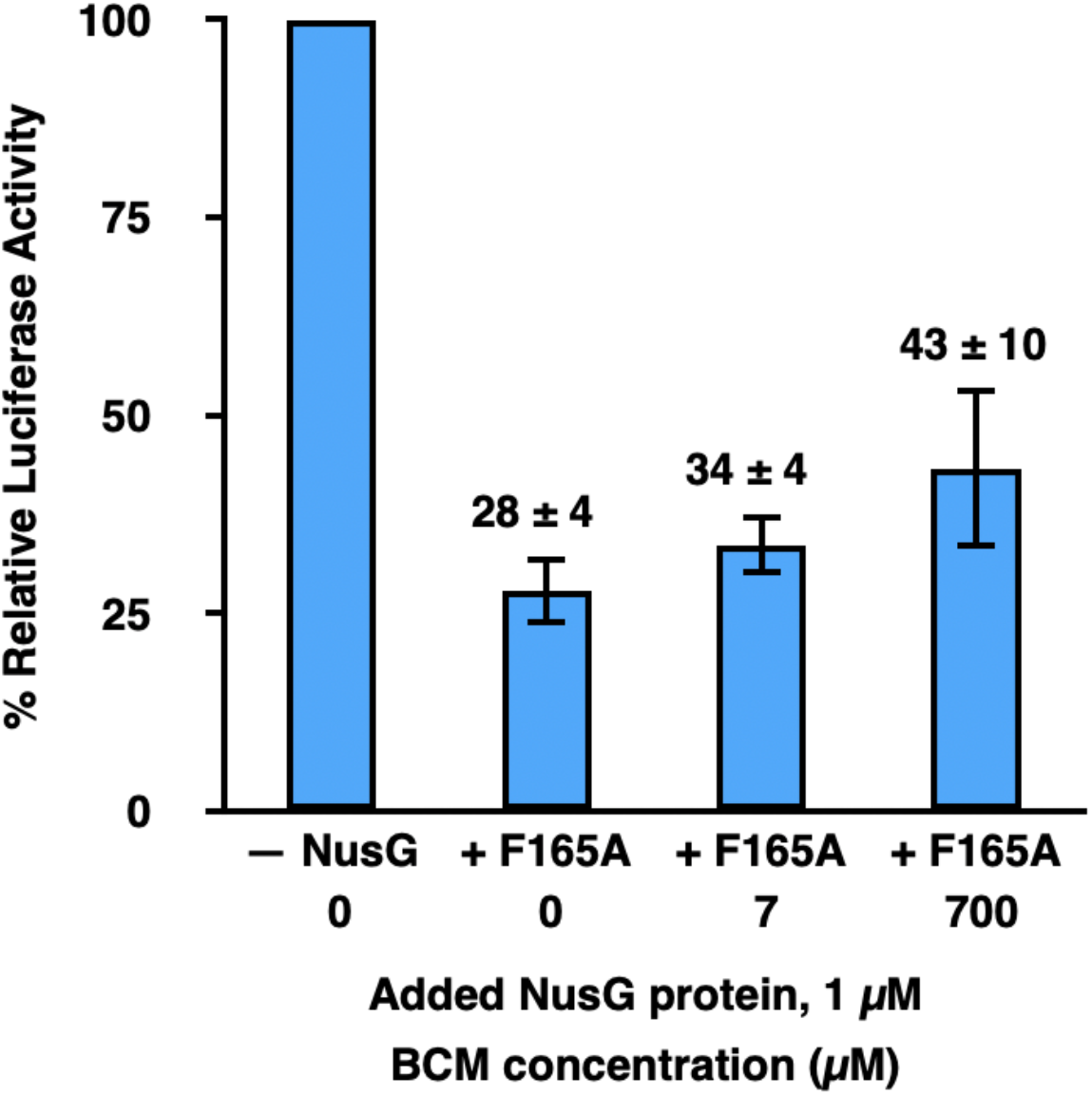
Prevention of premature, Rho-dependent transcription termination is not the primary mechanism through which NusG-mediated tethering of RNAP to the lead ribosome enhances gene expression in transcription-translation reactions. Bar graph plotting the % relative luciferase activity of transcription-translation reactions performed using standard extract in the absence of any added NusG proteins and 0 μM BCM (− NusG 0), in the presence of 1 μM added NusG-F165A and 0 (+ F165 0), 7 (+ F165 7), or 700 (+ F165 700) μM BCM. The luciferase activities in + F165 0, + F165 7, and + F165 700 are reported relative to the luciferase activity of – NusG, which is set to 100%. Error bars represent the standard deviations of each measurement.

We next asked if RNAP backtracking was responsible for the inhibition of luciferase mRNA synthesis by NusG-NTD or NusG-F165A. Accordingly, we performed transcription-translation reactions in an S30 extract generated from an *E. coli* MG1655-based strain carrying an RNAP that exhibits reduced backtracking. Specifically, this extract was generated from a strain harboring a substitution mutation in which the histidine at residue position 1244 of the β subunit of RNAP has been mutated to a glutamine (MG1655 *ΔintR-kilR::Cam*^*R*^ *rpoB*35*), hereafter referred to as the *rpoB*35* strain and extract. The H1244Q substitution mutation in the β subunit of RNAP has been previously shown to suppress RNAP backtracking [17, 18].

The results shown in Figure 3 demonstrate that transcription-translation reactions performed in *rpoB*35* extract are significantly more resistant to NusG-NTD- and NusG-F165A-mediated disruption of RNAP-ribosome coupling and consequent inhibition of transcription than transcription-translation reactions performed in standard extract. Specifically, the % relative luciferase activities measured for transcription-translation reactions performed in *rpoB*35* extract with 1 μM added NusG-NTD or NusG-F165A were 65 ± 10 % and 92 ± 17 %, respectively, compared to 34 ± 3 % and 28 ± 4 %, respectively, for the analogous reactions performed in standard extract. Furthermore, when NusG-NTD was titrated from 0.3–5 μM, transcription-translation reactions performed in *rpoB*35* extract were significantly more resistant to inhibition by NusG-NTD than those performed in standard extract.

**Figure 3.**
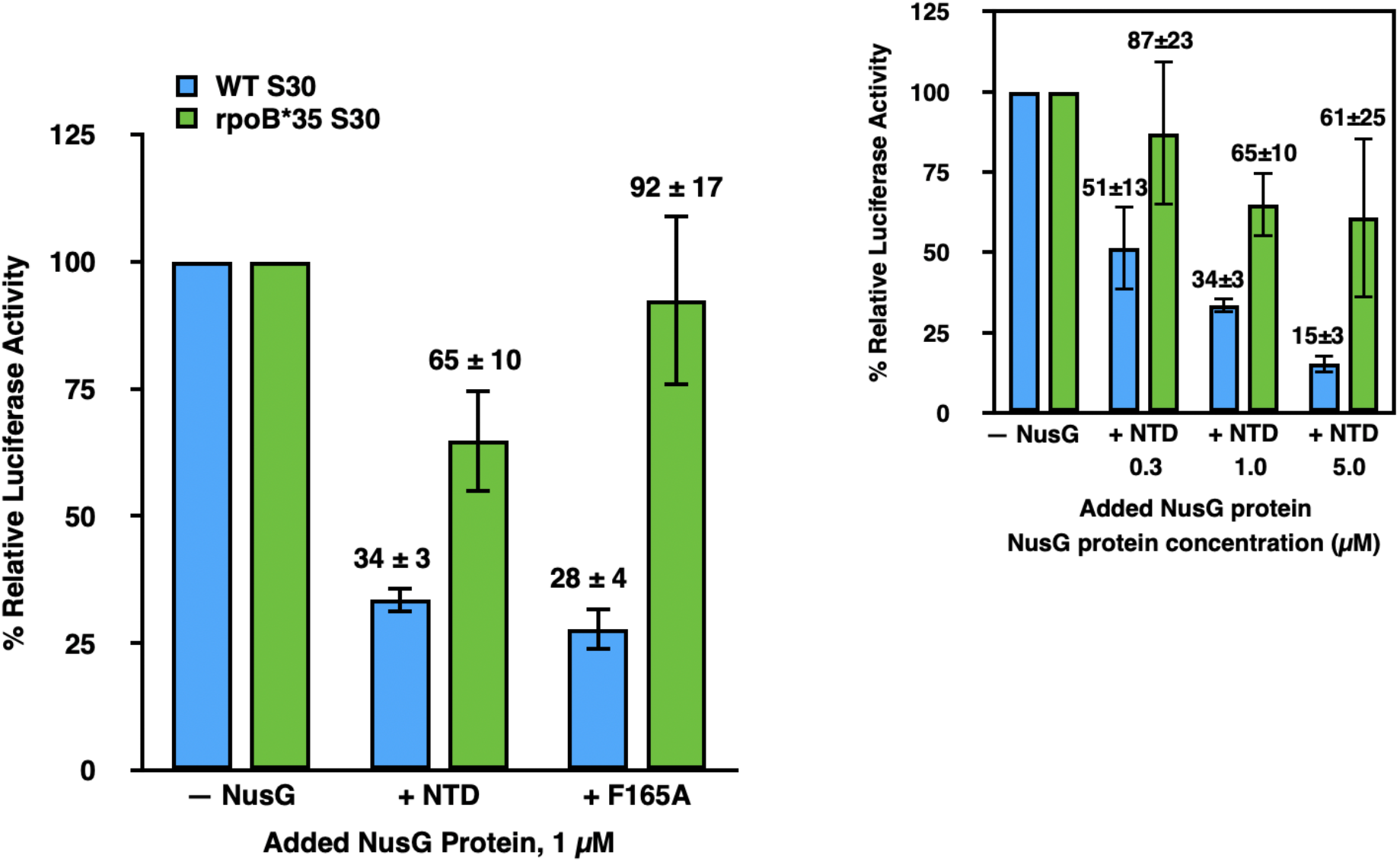
Suppression of RNAP backtracking during transcription is the primary mechanism though which NusG-mediated tethering of RNAP to the lead ribosome enhances gene expression in transcription-translation reactions. Bar graph plotting the % relative luciferase activity of transcription-translation reactions using performed using standard (blue bars) or *rpoB*35* (green bars) extracts in the absence of any added NusG proteins (− NusG) or in the presence of 1 μM added NusG-NTD (+ NTD) or NusG-F165A (+ F165A). For the standard and *rpoB*35* extracts, the luciferase activities in + NTD or + F165A are reported relative to the luciferase activities of – NusG in standard or *rpoB*35* extracts, respectively, which are each set to 100%. Error bars represent the standard deviations of each measurement. The inset is a bar graph plotting the % relative luciferase activity of transcription-translation reactions using standard or *rpoB*35* extracts in the absence of any added NusG proteins (– NusG) or in the presence of added NusG-NTD at 0.3 (+ NTD 0.3), 1 (+ NTD 1), or 5 (+ NTD 5) μM. For the standard and *rpoB*35* extracts, the luciferase activities of + NTD 0.3, + NTD 1, and + NTD 5 are reported relative to the luciferase activities of – NusG in standard or *rpoB*35* extracts, respectively, which are each set to 100%. Error bars represent the standard deviations of each measurement.

The striking restoration of luciferase activity by backtracking-resistant RNAP indicates that NusG-NTD- and NusG-F165A-mediated disruption of coupling between RNAP and the lead ribosome allows uncoupled RNAP to enter into a non-productive backtracked state. Uncoupled RNAP may elongate more rapidly than the translating lead ribosome and, in doing so, become prone to backtracking. We conclude that a critical role of NusG is to suppress RNAP backtracking by coupling RNAP to the lead ribosome.

Figure 4 presents a mechanistic model summarizing our findings. This model is consistent with previous studies by Proshkin *et al.* [1] and Dutta *et al.* [19] suggesting that translation by the lead ribosome exerts control over the rate of transcription by preventing RNAP from spontaneously backtracking. Significantly extending these studies, the data we present here strongly suggests that the lead ribosome prevents RNAP backtracking through a mechanism in which RNAP is physically tethered to the lead ribosome by NusG. Although Turtola and Belogurov have previously reported that NusG exhibits inherent backtracking suppression activity in an *in vitro*, transcription-only biochemical system composed of purified components [20], because their system lacked ribosomes and the associated translation components, whether and how NusG might suppress RNAP backtracking within the context of transcription-translation coupling had remained unexplored until the present work. We note that our model does not necessarily exclude the possibility of additional mechanisms through which the lead ribosome prevents RNAP backtracking, including mechanisms in which RNAP forms direct, non-NusG-mediated interactions with the lead ribosome, interactions that have been observed in recent RNAP-ribosome structures [9, 11], at least under certain conditions [7] or mechanisms relying on stochastic coupling of RNAP to the lead ribosome [21, 22].

**Figure 4.**
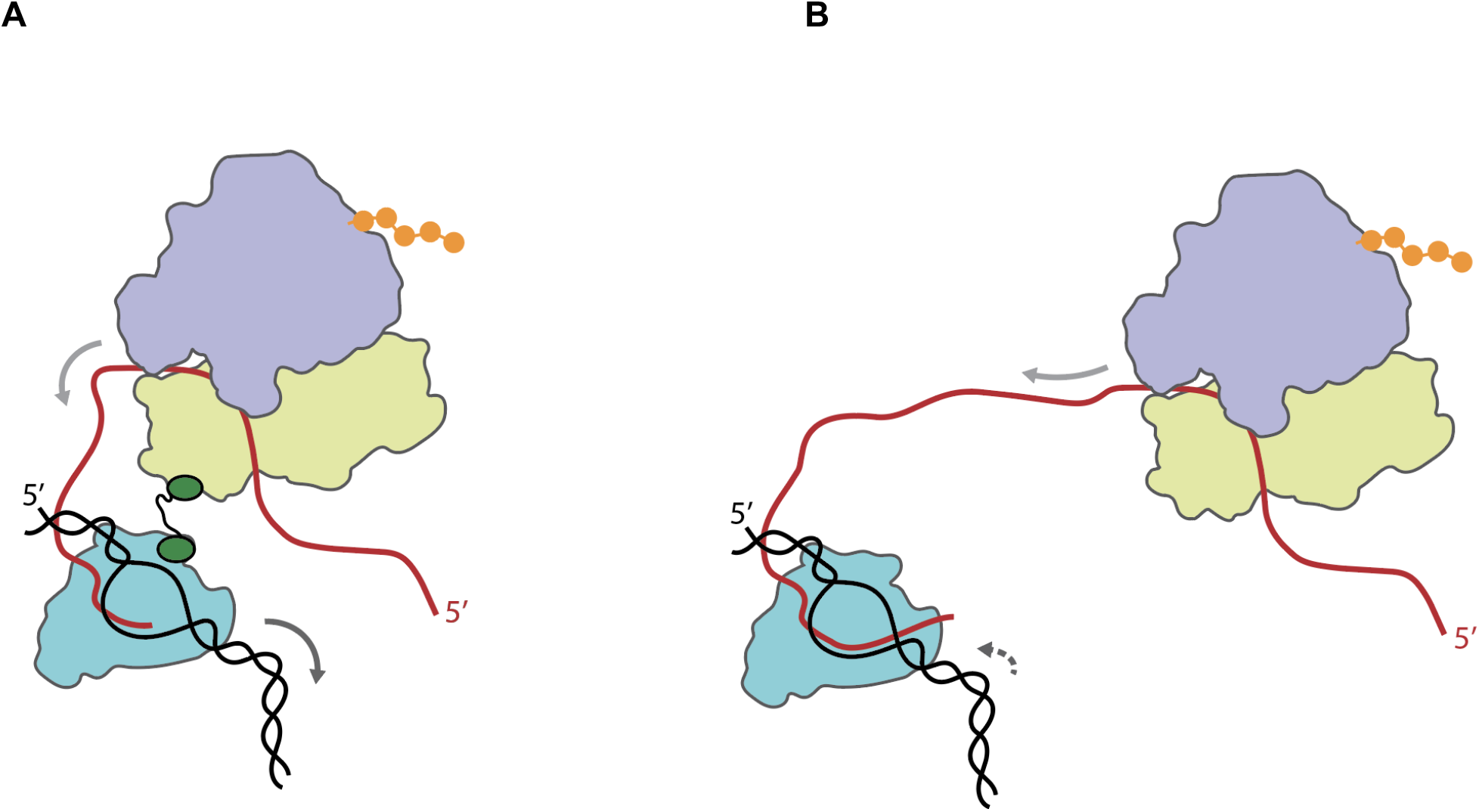
Mechanistic model showing how NusG-mediated tethering of RNAP to the lead ribosome increases the rate of transcription during transcription-translation coupling. **(A)** Transcription and translation in the presence of NusG (green)-mediated tethering of the transcribing RNAP (teal) to the lead ribosome (wheat and light blue for the small and large ribosomal subunits, respectively). NusG mediates tethering *via* interactions of the NusG CTD with the β’ and β subunits of RNAP and the NusG NTD with ribosomal protein uS10 within the small ribosomal subunit. Tethering enhances gene expression through a mechanism in which translation of the nascent mRNA (red) into protein (orange) by the NusG-tethered lead ribosome, the direction of which is shown by the solid light grey arrow over the mRNA, suppresses the tendency of RNAP to enter into a non-productive, backtracked state on the DNA template (black), thereby increasing the rate of transcription by RNAP, the direction of which is shown by the solid dark grey arrow over the DNA template. Tethering of RNAP to the lead ribosome may be further aided by NusA [7] not pictured. **(B)** Transcription and translation in the absence of NusG-mediated tethering of the transcribing RNAP to the lead ribosome. The lack of tethering enables RNAP to enter into the backtracked state, the direction of which is shown by the dashed dark grey arrow over the DNA template, thereby decreasing the rate of transcription by RNAP.

An advantage of conducting these studies in cellular extracts is that factors beyond NusG that might play a role in transcription-translation coupling, for example NusA [7], are included in the reactions. Consequently, straightforward extensions of the *in vitro* biochemical system described here should allow investigation of the role of such factors in the mechanism and regulation of transcription-translation coupling. Moreover, during gene expression *in vivo*, NusG is apparently recruited to a transcription elongation complex that is at some distance from its transcription promoter [23]. Here again, extension of the *in vitro* biochemical system we describe here should enable studies of the mechanism through which NusG is recruited to an elongating RNAP and through which NusG establishes interactions with the RNAP and the lead ribosome. Such studies should provide greater mechanistic insight into transcription-translation coupling, guiding the design of relevant structural constructs and prompting further studies into the structural basis of transcription-translation coupling.

## Materials and Methods

### Bacterial strains

The following strains were used in this study:

MG1655 *ΔintR-kilR::Cam*^*R*^[12]
MG1655 *ΔintR-kilR::Cam*^*R*^ *nusG::Kan*^*R*^
MG1655 *ΔintR-kilR::Cam*^*R*^ *rpoB*35*

The MG1655 wild-type-like strain, MG1655 Δ*intR-kilR::Cam*^*R*^, referred to herein as the wildtype MG1655 parent strain, is an MG1655 strain with an *intR-kilR* deletion that allows for the deletion of the otherwise essential gene, *nusG* [13]. Originally named RSW485, this strain was first used in Cardinale *et al.* 2008 and the Supplementary Information for Cardinale *et al.* 2008 therefore details the construction of the strain [12]. The two MG1655 strains used in the current study were constructed in the MG1655 Δ*intR-kilR::Cam*^*R*^ background. The strain herein referred to as *nusG* KO is MG1655 *ΔintR-kilR::Cam*^*R*^ *nusG::Kan*^*R*^. The knock-out of *nusG* was accomplished as described for the MDS42 *nusG::Kan*^*R*^, RSW422, in Cardinale *et al.* 2008 [12]. The strain herein referred to as *rpoB*35* is MG1655 Δ*intR-kilR::Cam*^*R*^ *rpoB*35*. The *rpoB*35* strain carries a codon mutation in rpoB, the gene which encodes for the β subunit of RNAP, that alters one amino acid residue (β H1244Q) [24, 25].

### Preparation of S30 cellular extracts

A culture of the *nusG* KO strain (to prepare *nusG* KO extract), wildtype MG1655 parent strain (to prepare standard extract), or the *rpoB*35* strain (to prepare *rpoB*35* extract) was grown in Terrific Broth with a 1% glucose supplement at 37 °C to an optical density at 600 nm (OD_600_) of 0.8-1.0 and the culture was subsequently cooled by placing in an ice bath for 1 hr. Cells were pelleted by centrifugation and washed in Extract Buffer (10 mM tris(hydroxymethyl)aminomethane (Tris) acetate (OAc) at a pH at 4 °C of 7.5, 1 mM dithiothreitol (DTT), 14 mM magnesium acetate (Mg(OAc)2), and 60 mM potassium chloride (KCl)). Cells were resuspended in 1 mL of Extract Buffer per 1g of wet cell weight. 250 μL Protease Inhibitor Cocktail (18 mM 4-(2-aminoethyl)benzenesulfonyl fluoride hydrochloride (AEBSF) (Sigma, No. A8456), 1.7 mM bestatin (Sigma, No. B8385), 290 μM pepstatin A (Sigma, No. P4265), and 220 μM E-64 (Sigma, No. E3132)) per 1g of wet cell weight and 1 Unit of 30 Units/μL RNase Inhibitor (from human placenta; New England Biolabs (NEB), No. M0307) per μl of total volume was added to the resuspended cell solution. Cells were lysed in a French press, 1 μL of 1 M DTT per ml of lysate was added to the lysate, and the lysate was gently mixed. The lysate was centrifuged at 30,000 ×g for 30 min at 4 °C, the supernatant was decanted into a fresh centrifuge bottle, and the supernatant was then centrifuged a second time at 30,000 ×g for 30 min at 4 °C. The supernatant was then transferred to a dialysis bag (molecular weight cutoff (MWCO) = 3.5 kDa) and dialyzed three times for 1 hr against 1 L of Extract Buffer at 4 ºC, replacing the used 1 L of buffer with a fresh 1 L of buffer between each time. The S30 extract was clarified by centrifugation one last time at 4,000 ×g for 10 min at 4 °C. To quantify the total concentration of biomolecules in the 30S extract, we measured the ultraviolet (UV) absorbance of the 30S extract at 280 nm (A_280_) and used A_280_ Units/μL as a proxy for the total concentration of biomolecules (the final *nusG* KO, standard, and *rpoB*35* extracts used in this study were 162 A_280_ Units/μL, 287 A_280_ Units/μL, and 171 A_280_ Units/μL, respectively). The S30 extract was then aliquoted, flash-frozen in liquid nitrogen, and stored at −80 °C until use.

### Luciferase-encoding plasmids and expression and purification of luciferase-encoding mRNA

The pBESTluc plasmid was obtained from Promega (No. L1020) and contains the eukaryotic firefly luciferase gene positioned downstream from a Ptac promoter and a ribosome binding site [26]. Notably, the luciferase gene encoded by the pBESTluc plasmid lacks an N-utilization (*nut*) site and therefore does not promote the assembly of an RNAP anti-termination complex. We constructed the pBESTlucT7 plasmid by replacing the Ptac promoter in the pBESTluc plasmid with the T7 RNAP promoter. The pBESTluc and pBESTlucT7 plasmids used in this study were electroporated into *E. coli* XL1-Blue (Agilent) and 10G (Lucigen) electrocompetent cells, respectively, and purified using the QIAprep Spin Miniprep Kit (No. 27104) or QIAGEN HiSpeed Plasmid Maxi Kit (No. 12663), depending on desired scale of yield. The concentration of the resulting pBESTluc plasmid solution, in μg/μL, was calculated from the A_260_, as measured using a NanoDrop spectrophotometer (Thermo Fisher Scientific), and an extinction coefficient of 0.020 (μg/ml)^−1^ cm^−1^. To perform transcription-translation reactions, we added purified pBESTluc plasmid directly to transcription-translation reactions (*vide infra*). To perform translation-only reactions, we first used the HiScribeTM T7 Quick High Yield RNA Synthesis Kit (NEB, No. E2050S) to *in vitro* transcribe the pBESTlucT7 plasmid and generate pBESTlucT7 mRNA using the protocol provided in the manufacturer’s instruction manual [27]. Upon completion of transcription, the reaction was treated with DNase I (NEB, No. M0303S) to degrade the pBESTlucT7 plasmid. DNase I was then inactivated by adding 2 μl of 0.2 M ethylenediaminetetraacetic acid (EDTA) per 20 μl T7 transcription reaction and heating at 70 °C for 10 min. The completeness of the DNA template degradation was confirmed by 5% denaturing polyacrylamide gel electrophoresis (D-PAGE). Owing to difficulties in further mRNA purification, to perform translation-only reactions, we added pBESTluc mRNA, as a standard amount of the T7 RNA transcription reaction product, directly to translation-only reactions (*vide infra*).

### Over-expression and purification of NusG-WT, NusG-NTD, and NusG-F165A proteins

Purified NusG-WT, NusG-NTD, and NusG-F165A proteins were a generous gift from Prof. Paul Rösch at the University of Bayreuth. Over-expression and purification of NusG-WT, NusG-NTD, and NusG-F165A proteins is described in Burmann *et al*. 2011 [14]. Purified NusG-WT and NusG-F165A were stored in a storage buffer composed of 10 mM Tris hydrochloride (Tris-HCl) at a pH at room temperature (~23 °C) of 7.5 and 150 mM sodium chloride (NaCl). Purified NusG-NTD was stored in a storage buffer of 50 mM Tris-HCl at a pH at room temperature (~23 °C) of 7.5 and 150 mM NaCl.

Using solution NMR spectroscopy experiments [2, 14] and *in vitro* transcription assays [5], we have previously validated the proper folding and expected biochemical activities of these purified NusG-WT, NusG-NTD, and NusG-F165A proteins. Moreover, we have shown that NusG-NTD is toxic when expressed in *E. coli* strains containing endogenous levels of NusG-WT [3], suggesting that NusG-NTD, and, by extension, NusG-F165A, can bind RNAP in a manner that competes with NusG-WT. Collectively, these observations support the design of our transcription-translation and translation-only assays, in which we add excess concentrations of purified NusG-WT, NusG-NTD, or NusG-F165A to standard extract and assess how these proteins compete with the endogenous NusG-WT that is found in standard extract.

### Transcription-translation reactions, translation-only reactions, luciferase activity assays, and data analyses

Transcription-translation reactions were performed by combining in an Eppendorf tube *v*_DNA_ μL of a pBESTluc plasmid DNA solution, where *v*_DNA_ μL is the volume of a pBESTluc plasmid solution at a particular μg/μL concentration that is required to deliver 2 μg of pBESTluc plasmid DNA to the reaction; 5 μL of a solution that is 1 mM in each of the 20 essential amino acids (Promega, No. L4461); 20 μl of Promega S30 Premix without Amino Acids (No. L512A-C); *v*_S30_ μL of S30 cell extract, where *v*_S30_ μL is the volume of S30 extract at a particular A_280_ Units/μL concentration that is required to deliver 2,000 A_280_ Units of S30 extract to the reaction; and *v*_H2O_ μL Nanopure water (H_2_O), where *v*_H2O_ μL is the volume of Nanopure H_2_O that is required to achieve a final reaction volume of 50 μL. Transcription-translation reactions were incubated for 60 minutes at 37 °C and subsequently stopped by incubating on ice for 5 minutes. The transcription-translation reaction was then shifted to room temperature (~23 °C); 50 μL of Promega Steady-Glo Luciferase Assay Reagent (No. E2520) was added to the 50 μL transcription-translation reaction; 20 μL of the resulting 100 μL Luciferase Assay Reagent-containing reaction mixture was transferred to a white, flat-bottom 96-well plate; the plate was incubated for 10 min at room temperature (~23 °C); and, immediately following the 10 min incubation, the luminescence was quantified in relative light units (RLU) using a Tecan Infinite 200 Multimode Plate Reader.

Translation-only reactions were performed in a manner identical to that of transcription-translation reactions with two exceptions. The first exception was that the *v*_DNA_ μL of the pBESTluc plasmid solution was replaced by 6 μL of a DNase I-treated T7 transcription reaction solution (*vide supra*) such that pBESTlucT7 mRNA could be delivered to the reaction. We accounted for slight variations in the mRNA concentration of individual DNase I-treated T7 transcription reaction solutions by using a single DNase I-treated T7 transcription reaction solution to perform multiple translation-only reactions in parallel and always including a ‘reference’ translation-only reaction within each group of the parallelized translation-only reactions, as described in the next paragraph. The second exception was that the volume of the 100 μL Luciferase Assay Reagent-containing reaction mixture that was transferred to the 96-well plate was increased from 20 μl to 80 μl in order to make up for the fact that translation-only reactions generate less luciferase and, correspondingly, lower RLU than transcription-translation reactions. Because our experiments and analyses make use of a ‘reference’ reaction, as described in the following paragraph, it was unnecessary to correct the data for this difference in the volume of Luciferase Assay Reagent-containing reaction mixture that was transferred to the plates for the transcription-translation and translation-only reactions.

To account for possible preparation-to-preparation, experiment-to-experiment, and/or day-to-day variations in the concentrations or activities of reaction components, our ability to reproducibly assemble the reactions, and/or the performance of equipment and instruments and enable comparison of our results across multiple reaction component preparations, experiments, and days, we always performed multiple reactions in parallel, in groups of up to 12 reactions, and consistently included corresponding ‘reference’ reactions within each group of parallelized reactions. For transcription-translation reactions performed in standard extract, the corresponding reference reaction was a transcription-translation reaction performed in standard extract in the absence of added NusG proteins and BCM. Analogously, for translation-only reactions performed in standard extract, the corresponding reference reactions was a translation-only reaction performed in standard extract in the absence of added NusG proteins. For transcription-translation reactions performed in *rpoB*35* extract, the corresponding reference reaction was a transcription-translation reaction performed in *rpoB*35* extract in the absence of added NusG proteins. Having defined these reference reactions, the % relative luciferase activity of each transcription-translation or translation-only reaction performed within a group of parallelized reactions could be calculated as [(RLU of the transcription-translation or translation-only reaction performed within a group of parallelized reactions in standard or *rpoB*35* extract, in the absence or presence of added NusG proteins, and/or in the absence or presence of BCM within one group of parallelized reactions) / (RLU of the corresponding reference reaction performed within the same group of parallelized reactions) × 100]. % relative luciferase activities calculated in this manner account for possible variations in the concentrations or activities of assay components, our ability to reproducibly assemble the reactions, and/or the performance of equipment and instruments and can therefore be compared across multiple assay component preparations, experiments, and days.

Two technical replicates were performed for the translation-only reactions executed in standard extract in the absence of any added NusG proteins and in the presence of 1 μM added NusG-WT, NusG-NTD, and NusG-F165A and for the transcription-translation reactions executed in *rpoB*35* extract in the presence of NusG-NTD at 5 μM. A minimum of three technical replicates were performed for all other reactions. Replicates were used to calculate the mean and the standard deviation of the % relative luciferase activity.

## Acknowledgements

We would like to thank Dr. Robert S. Washburn for constructing the *nusG* KO, wildtype MG1655 parent, and *rpoB*35* strains and Dr. Martin Strauβ for performing β-galactosidase-based, *in vivo* cell biology experiments. The purified NusG-WT, NusG-NTD, and NusG-F165A proteins used in this study were a generous gift from Prof. Paul Röche at the University of Bayreuth. This work was supported by funds to R.L.G. from the National Institutes of Health (NIH) (R01 GM 084288 and R01 GM 137608) and M.E.G. from NIH (R01 GM 37219). E.J.B was supported by a National Science Foundation Graduate Research Fellowship (DGE 1644869).

## CRediT Authorship Contribution Statement

**Elizabeth J. Bailey:** Conceptualization, Methodology, Resources, Investigation, Validation, Formal Analysis, Data Curation, Writing - original Draft, Writing - Review & Editing, Visualization, Funding Acquisition. **Max E. Gottesman:** Conceptualization, Methodology, Resources, Writing - original Draft, Writing - Review & Editing, Visualization, Supervision, Project Administration. **Ruben L. Gonzalez, Jr.:**Conceptualization, Methodology, Resources, Data Curation, Writing - Review & Editing, Visualization, Supervision, Project Administration, Funding Acquisition.

