## Supplementary Information for "NusG links transcription and translation in *Escherichia coli* extracts"

### Supplemental Materials and Methods

#### *Bacterial strains and plasmid*

The following MDS42-based strains and plasmid were used in the *in vivo* β-galactosidase assays designed to test whether expression of a plasmid-encoded copy of NusG-WT can complement the lack of NusG in a *nusG* deletion strain (Supplemental Figure S1):

MDS42 [1]

MDS42, transformed with pRM431, a wildtype NusG-encoding plasmid [2]

MDS42 *nusG::Kan*$^R$ [3]

MDS42 *nusG::Kan*$^R$, transformed with pRM431 [2]

The MDS42 strain [1] is a previously described, reduced-genome variant of the MG1655 strain [4]. The pRM431 plasmid is a previously described wildtype NusG-encoding plasmid in which *nusG* is under the control of the p*trc* promoter. The MDS42 *nusG::Kan*$^R$ [3] strain is a previously described MDS42-based *nusG*-deletion strain that was formed by replacing *nusG* with a kanamycin resistance cassette in MDS42. Interestingly, S30 extracts prepared from MDS42 did not exhibit detectable luciferase activity in *in vitro* transcription-translation reactions (data not shown). Instead, S30 extracts exhibiting robust luciferase activity in *in vitro* transcription-translation reactions could only be prepared from the MG1655-based strains described in the main text and figures of this article. This is an intriguing result in that MDS42 is derived from an MG1655 strain in which a genomic segment representing ~15% of the genome and composed of non-essential genes, including insertion sequences, prophages, and pseudogenes, was deleted [1]. A possible explanation for this observation is that a gene essential for *in vitro* S30 extract activity has been deleted in MDS42. Alternatively, a gene whose function is inhibitory in these extracts is expressed in both strains, but is suppressed in MG1655 and not in MDS42. Regardless of the explanation, however, these initial findings drove us to use MG1655 as the background strain for preparing the S30 extracts that we subsequently employed in our transcription-translation and translation-only reactions and all of the other experiments in this study.

***β-galactosidase assays and data analysis***

β-galactosidase assays were performed in four strain conditions: (*i*) the MDS42 strain (MDS42(WT)), (*ii*) the MDS42 strain transformed with the pRM431 wildtype NusG-encoding plasmid (MDS42(WT) + NusG), (*iii*) the MDS42 *nusG::Kan*$^{R}$ *nusG*-deletion strain (MDS42(*nusG* KO)), and (*iv*) the MDS42 *nusG::Kan*$^{R}$ *nusG*-deletion strain transformed with the pRM431 wildtype NusG-encoding plasmid (MDS42(*nusG* KO) + NusG). For each of four biological replicates performed at each strain condition, cells were grown in Luria-Bertani (LB) liquid culture at 37 °C and, at mid-log phase, isopropyl β-d-1-thiogalactopyranoside (IPTG) was added to the culture to a final concentration of 1 mM to induce β-galactosidase and NusG expression. Four hours after induction, the β-galactosidase activity was assayed and Miller unit value was calculated by following the OpenWetWare protocol published by Moore [5], which is based on an earlier publication by Zhang and Bremer [6]. Miller unit values for the four replicates at each strain condition were averaged and the standard deviation was calculated at each strain condition. Results are plotted as % relative β-galactosidase activity in which the activity of β-galactosidase at each condition is reported relative to MDS42(WT), which is set to 100% (Figure S1).

***Western blots***

Western blots were performed for standard and *nusG* KO extracts using primary antibodies against the β' subunit of RNAP (Neoclone, No. NT73) and ribosomal protein S3 (Developmental Studies Hybridoma Bank, University of Iowa, No. 373C9C3A1) on a 0.2 μm nitrocellulose membrane. Briefly, samples were mixed with 1/4 volume of 4× NuPAGE LDS Sample Buffer (Invitrogen, No. NP0007); heat-denatured at 100 °C for 2 min; and electrophoresed on a

NuPAGE bis-(2-hydroxyethyl)-amino-tris(hydroxymethyl)-methane (Bis-Tris; free base), SDS, 4-12% polyacrylamide gradient gel (Invitrogen, No. NP0321BOX) using NuPAGE 2-(N-morpholino)ethanesulfonic acid (MES) SDS Running Buffer (Invitrogen, No. NP0002). Transfer to a 0.2 µm nitrocellulose membrane was performed in Transfer Buffer (25 mM *N*,*N*-Bis(2-hydroxyethyl)glycine (Bicine), 25 mM Bis-Tris (free base), 1 mM EDTA, and pH at room temperature (~23 °C) of 7.2) at 100 V for 1 hr. Membranes were soaked in Blocking Buffer (5% w/v nonfat dry milk dissolved in 1× Phosphate-Buffered Saline with 0.1% Tween (PBST) Buffer (10 mM dibasic sodium phosphate ($Na_2HPO_4$), 1.8 mM monobasic potassium phosphate ($KH_2PO_4$), 137 mM sodium chloride (NaCl), 2.7 mM potassium chloride (KCl), and 0.1% w/v Tween-20 detergent) for 15 min, then incubated with primary antibodies overnight. Prior to using, primary antibodies were diluted 1,000-fold using 0.5× PBST Buffer. Blots were washed in 1× PBST Buffer and then incubated with the secondary antibody for 1 hr. The secondary antibody was IRDye 800CW-labeled goat anti-mouse immunoglobulin G (IgG) (926-32210, Li-Cor). Prior to using, the secondary antibody was diluted 3,000-fold using a 1:1 mixture of 1× PBST Buffer and PBS Buffer (10 mM $Na_2HPO_4$, 1.8 mM $KH_2PO_4$, 137 mM NaCl, and 2.7 mM KCl). Blots were then washed three times in 1× PBST Buffer for 5 min each time and with a fresh change of buffer between washes and subsequently washed a final time in PBS Buffer for 5 min prior to imaging with a Li-Cor Odyssey imaging system. The results of the Western blots are consistent with our conclusion that equivalent ultraviolet (UV) absorbance at 280 nm ($A_{280}$) Units of each S30 extract contain approximately equivalent amounts of RNAP and ribosomes (Figure S2). These findings support our experimental design, in which we use equivalent $A_{280}$ Units of each S30 extract as a way of matching the concentrations of RNAP, ribosomes, and other key cellular protein components across S30 extracts prepared at different times and/or from different strains.

***RNA-Seq and analysis***

Wildtype MG1655 parent and *nusG* KO strains were grown as we described for the preparation of S30 cellular extracts (*i.e.*, in Terrific Broth (TB) with a 1% glucose supplement to an optical density at 600 nm ($OD_{600}$) of 0.8-1.0; Materials and Methods). Cultures were cooled in an ice bath for 1 hr. Cells were pelleted by centrifugation and washed in Extract Buffer (Materials and Methods). RNA extraction, next-generation sequencing, and data analysis was performed by Girihlet, Inc. (Oakland, CA).

The Girihlet data analysis pipeline that was used for the analysis is described in reference [7]. Briefly, the number of reads mapping to each gene (counts per gene) was first determined. The counts per gene were then normalized to the sequence length of the corresponding genes. Subsequently, quantile normalization was used to make the distributions of the normalized counts per gene from the wildtype MG1655 parent and *nusG* KO strains equivalent. $Log_2$ fold-changes between the normalized counts per gene for each gene of the wildtype MG1655 parent and *nusG* KO strains were then calculated as $\log_2$(*nusG* KO – $m$) – $\log_2$(wildtype MG1655 parent – $m$), where $m$ is the median of all the normalized counts per gene for a given gene.

**Supplementary Figures**

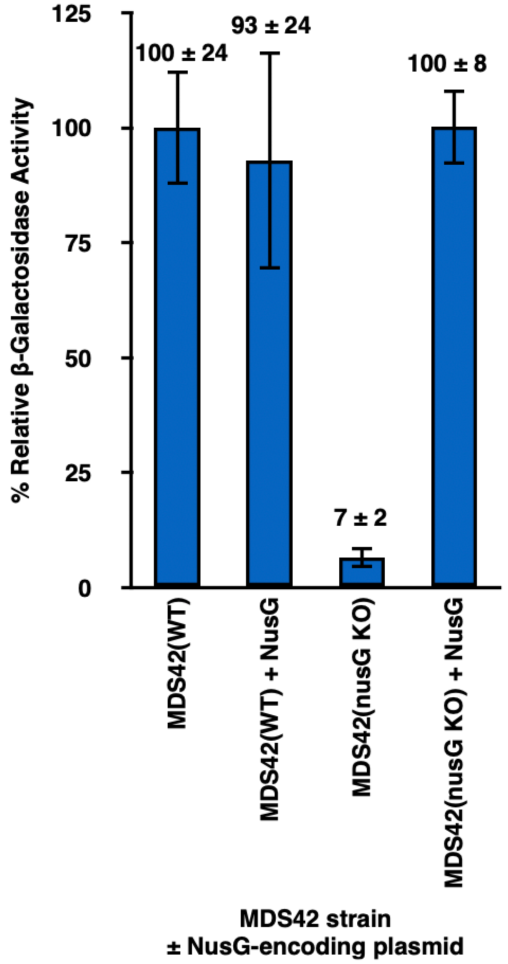


**Figure S1. The β-galactosidase expression defect exhibited by the MDS42(*nusG* KO) strain can be fully complemented by expression of wildtype NusG.** Bar graph plotting the % relative β-galactosidase activity observed in the *in vivo* β-galactosidase assays performed at the MDS42(WT), MDS42(WT) + NusG, MDS42(*nusG* KO), and MDS42(*nusG* KO) + NusG strain conditions. The β-galactosidase activities at the MDS42(WT) + NusG, MDS42(*nusG* KO), and MDS42(*nusG* KO) + NusG strain conditions are reported relative to the β-galactosidase activity at the MDS42(WT) strain condition, which is set to 100%. Error bars represent the standard deviations of four replicates for each condition.

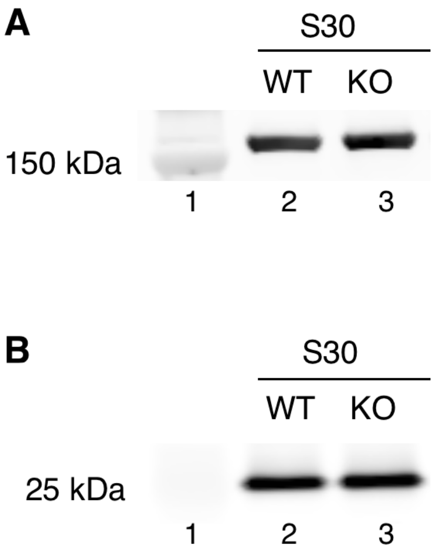


**Figure S2. Western blots demonstrate that S30 extracts prepared from the wildtype MG1655 parent and *nusG* KO strains contain similar RNAP and ribosome concentrations.** Western blots of S30 extracts prepared from the wildtype MG1655 parent and *nusG* KO strains were performed with primary antibodies raised against **(A)** the β’ subunit of RNAP and **(B)** ribosomal protein S3. In each panel, Lane 1 is a molecular weight standard, Lane 2 is the S30 extract from the wildtype MG1655 parent strain (WT), and Lane 3 is the S30 extract from the *nusG* KO strain (KO).

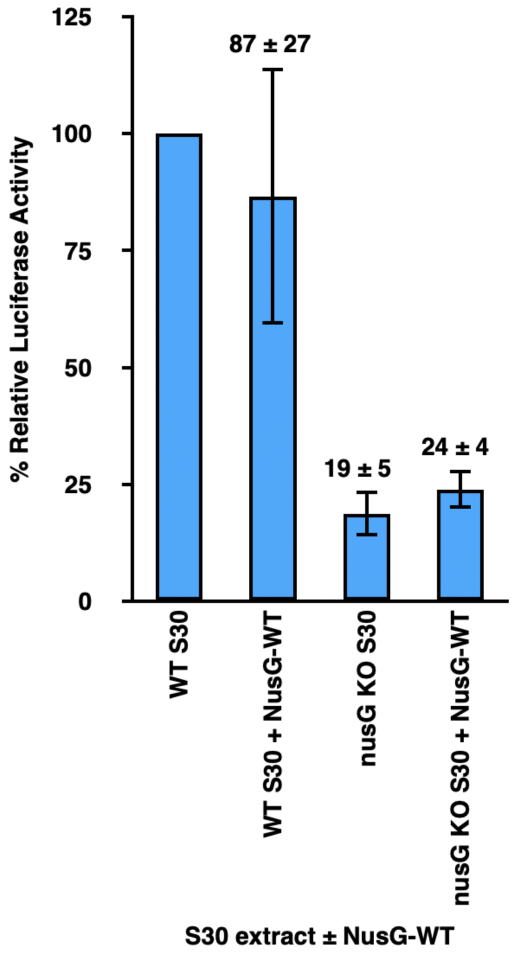


**Figure S3. The luciferase expression defect exhibited by the S30 extract prepared from the *nusG* KO strain cannot be complemented by the addition of NusG-WT.** Bar graph plotting the % relative luciferase activity of transcription-translation reactions performed using wildtype MG1655 parent (WT S30) or *nusG* KO (*nusG* KO S30) extracts in the absence of any added NusG proteins or in the presence of 5 μM added NusG-WT (WT S30 + NusG-WT and *nusG* KO + NusG-WT, respectively). The luciferase activities of *nusG* KO S30, WT S30 + NusG-WT, and *nusG* KO + NusG-WT are reported relative to the luciferase activity of WT S30, which is set to 100%. Error bars represent the standard deviations of each measurement.

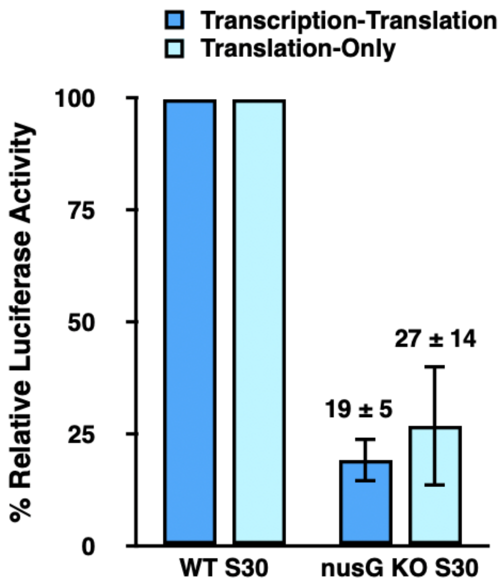


**Figure S4. Translation-only reactions performed in S30 extracts prepared from the *nusG* KO exhibit luciferase activities that are as extremely low as those exhibited by transcription-translation reactions performed in the same S30 extracts, a result consistent with the notion that this S30 extract harbors pleiotropic defects in translation.** Bar graph plotting the % relative luciferase activity of transcription-translation (dark blue bars) and translation-only (light blue bars) reactions performed using S30 extracts prepared from the wildtype MG1655 parent (WT S30) or *nusG* KO (*nusG* KO S30) strains. The luciferase activities of the transcription-translation and translation-only reactions in *nusG* KO S30 are reported relative to those of the transcription-translation or translation-only reactions, respectively, in WT S30, which are each set to 100%. Error bars represent the standard deviations of each measurement.

### Supplementary Tables

**Table S1. Translation-associated genes exhibiting the largest $\log_2$ fold-changes in expression between the wildtype MG1655 parent and *nusG* KO strains.** Analysis of RNA-Seq data from wildtype MG1655 parent and *nusG* KO strains reveals significant up- and down-regulation of a number of translation-associated genes. The difference in expression between wildtype MG1655 parent and *nusG* KO strains is calculated as $\log_2(nusG\ \mathrm{KO} - m) - \log_2(\mathrm{wildtype\ MG1655\ parent} - m)$, where $m$ is the median of all the normalized counts per gene for a given gene.

| Gene | $\log_2$ fold-change | Function[1] |
|---|---|---|
| serW | –2.266 | tRNA-Ser(GGA) |
| aspV | –2.193 | tRNA-Asp(GUC) |
| trpT | –2.162 | tRNA-Trp(CCA) |
| serT | –2.108 | tRNA-Ser(UGA) |
| rrfF | –2.105 | 5S ribosomal RNA |
| proM | –2.014 | tRNA-Pro(UGG) |
| pheV | –1.85 | tRNA-Phe(GAA) |
| thrV | –1.794 | tRNA-Thr(GGU) |
| aspU | –1.618 | tRNA-Asp(GUC) |
| valU | –1.601 | tRNA-Val(UAC) |
| rpmJ | 1.481 | 50S ribosomal subunit protein L36 |
| valX | –1.345 | tRNA-Val(UAC) |
| selC | –1.298 | tRNA-Sec(UCA) |
| sra | –1.246 | 30S ribosomal subunit protein S22 |
| thrW | –1.233 | tRNA-Thr(CGU) |
| rrfA | –1.219 | 5S ribosomal RNA |
| infA | –1.124 | translation initiation factor IF-1 |
| argU | –1.098 | tRNA-Arg(UCU) |
| valY | –1.073 | tRNA-Val(UAC) |
| glyX | –1.055 | tRNA-Gly(GCC) |
| rplO | 1.054 | 50S ribosomal subunit protein L15 |
| ileV | –1.045 | tRNA-Ile(GAU) |

[1] Based on functions identified in the EcoCyc database (https://ecocyc.org)

**Table S2. Additional notable genes exhibiting the largest $log_2$ fold-changes in expression between the wildtype MG1655 parent and *nusG* KO strains.** Analysis of RNA-Seq data from wildtype MG1655 parent and *nusG* KO strains reveals significant up- and down-regulation of a number of notable genes. The difference in expression between wildtype MG1655 parent and *nusG* KO strains is calculated as $\log_2(nusG\text{ KO} - m) - \log_2(\text{wildtype MG1655 parent} - m)$, where *m* is the median of all the normalized counts for a given RNA.

| Gene | $log_2$ fold-change | Function[1] |
|---|---|---|
| ffs | –2.746 | 4.5S RNA component of the signal recognition particle (binds ribosomes) |
| fnrS | –2.382 | FnrS small regulatory RNA |
| csrC | –2.34 | RNA inhibitor of CsrA |
| csrB | –2.046 | CsrB small regulatory RNA |
| glmZ | –1.766 | small regulatory RNA GlmZ |
| rdlD | –1.72 | antisense regulatory RNA RdlD |
| sibD | –1.631 | small RNA SibD |
| sibE | –1.602 | small RNA SibE |
| glmY | –1.52 | small regulatory RNA GlmY |
| cysK | 1.304 | cysteine synthase A (a.a. biosynthesis) |
| ilvE | 1.241 | branched-chain-amino-acid aminotransferase (a.a. biosynthesis) |
| ilvD | 1.203 | dihydroxy-acid dehydratase (a.a. biosynthesis) |
| leuC | 1.169 | 3-isopropylmalate dehydratase subunit LeuC (a.a. biosynthesis) |
| hisL | –1.149 | his operon leader peptide (a.a. biosynthesis) |
| map | 1.127 | methionine aminopeptidase (binds ribosome exit tunnel) |
| leuA | 1.104 | 2-isopropylmalate synthase (a.a. biosynthesis) |
| leuD | 1.098 | 3-isopropylmalate dehydratase subunit LeuD (a.a. biosynthesis) |
| leuB | 1.089 | 3-isopropylmalate dehydrogenase (a.a. biosynthesis) |
| argG | –1.045 | argininosuccinate synthetase (a.a. biosynthesis) |
| dtd | –1.019 | D-aminoacyl-tRNA deacylase (deacylates D-aminoacyl-tRNAs) |
| tusD | –1.016 | sulfurtransferase complex subunit TusD (Δ*tusD* lacks the 2-thio modification of mnm5s2U in tRNA) |
| ilvL | –1.011 | ilvXGMEDA operon leader peptide (a.a. biosynthesis) |

[1] Based on functions identified in the EcoCyc database (https://ecocyc.org)
